# Scale-independent glide energetics in odontocete cetaceans

**DOI:** 10.64898/2026.06.29.735419

**Authors:** Vadim Pavlov, Teresa Salomone, Beverley McKeon

**Affiliations:** Stanford University, Hopkins Marine Station, Pacific Grove, USA; Stanford University, Department of Mechanical Engineering, Palo Alto, USA

**Keywords:** burst-and-glide swimming, cetacean energetics, glide optimization, body-form scaling, odontocetes, hydrodynamic drag, cost of transport

## Abstract

Cetaceans reduce the net cost of sustained swimming through intermittent locomotion, alternating active fluking with unpowered gliding. The energy balance of this strategy is central to understanding survival rates, population sustainability, and the effects of anthropogenic and environmental pressures. While active-phase energetics have been characterized extensively, the glide phase remains largely unexplored. Here we derive the optimal glide duration (*T*_opt_) and the maximum glide duration beyond which energy savings vanish (*T*_zero_) for three odontocetes spanning a 20-fold range in body mass, using high-fidelity CAD models and wall-modeled large eddy simulations. We show analytically that speed retention at *T*_opt_ and mass-specific peak energy savings are both fully determined by the active-to-passive drag ratio, propulsive efficiency, and swimming speed, independently of body morphometry and drag coefficient, and are therefore invariant across species at any given speed. These passive-phase optima extend the known size-independent active-phase invariants to the glide phase, towards a scale-independent energetic framework for burst-and-glide locomotion in small cetaceans.

## 1. Introduction

Sustained aquatic locomotion imposes high energetic costs on air-breathing vertebrates (Williams, 1999). The mass-specific cost of transport (COT) scales inversely and steeply with body size, so that small species must devote a substantially greater proportion of their energy budget to locomotion than large ones (Schmidt-Nielsen, 1972; Watanabe et al., 2011; Williams, 2022; Glarou et al., 2025). The present study focuses on transit swimming, the locomotion mode relevant to routine travel and migration, in which COT minimization is a relevant energetic objective. Burst-and-glide swimming alternates an active fluking phase with a passive glide phase and is a key kinematic mechanism by which cetaceans reduce net locomotor expenditure below that of continuous swimming at equivalent mean speed (Weihs, 1974; Fish, 1993; Liu et al., 2025a). The energetic advantage of this strategy arises from the difference in drag between active and passive phases, where total drag combines a pressure component and a viscous friction component. Active fluking destabilizes the boundary layer and modifies the wake, raising the friction component in particular, whereas cessation of fluking during a glide lowers both components and reduces the power required to maintain speed (Lighthill, 1971; Fish and Lauder, 2006; Borazjani and Sotiropoulos, 2008).

This drag difference is quantified by the ratio of active to passive drag, commonly denoted *g*. Empirical estimates of *g* in odontocetes span a wide range (approximately 3.2 to 10.6; Fish, 1993; Fish et al., 2014), with Webb (1975) reporting values of 6.3 and 9.4 for the long-beaked common dolphin and Dall’s porpoise respectively. Higher values at lower swimming speeds reflect the turbulence introduced by active fluke strokes when the boundary layer would otherwise remain partially laminar and the wake body-aligned. Previous studies have identified the former as the dominant effect within aquatic locomotion models. As speed increases and the boundary layer transitions toward turbulence even during passive gliding, the marginal turbulence penalty of active fluking diminishes and *g* declines (Lighthill, 1971; Fish, 1998).

Despite this established foundation, the energetic consequences for glide-phase optimization, and in particular how the optimal glide duration *T*_opt_ scales across species of widely differing body mass, remain unresolved. Bio-logging studies have documented considerable interspecific variation in observed glide durations and glide fractions (Noren et al., 2006; Williams et al., 2017; Arranz et al., 2019), yet mechanistic frameworks capable of attributing this variation to underlying energetic optima are scarce. Li et al. (2023) applied a hybrid computational fluid dynamics approach to fish as a model system and identified the cost-of-transport-minimizing coast duration through numerical parameter search. Zhu et al. (2025) performed an analogous analysis using high-fidelity simulations for undulatory swimmers. Both studies recovered results directionally consistent with *T*_opt_ but without a closed-form expression, and neither established *T*_zero_, the maximum glide duration beyond which cumulative savings become negative. Paoletti and Mahadevan (2014) approached this analytically, showing that the ratio of terminal speed to energy expenditure is independent of glide duration in their minimal model, but this result does not yield an optimal glide duration or a zero-savings threshold. The present framework resolves this by incorporating the active drag factor *g* into a closed-form derivation of *T*_opt_ and *T*_zero_, grounded in measurable hydrodynamic parameters. A key question that follows from the body-size scaling of *T*_opt_ is whether speed retention at the end of an optimal glide also scales with mass or is conserved across species. This quantity directly sets the kinetic energy deficit that the next stroke cycle must replenish and, if size-independent, would provide a unified benchmark for interpreting bio-logging data across species.

Here we derive, from first principles, *T*_opt_ and *T*_zero_ for three odontocete species spanning a 20-fold range in body mass, the false killer whale (*Pseudorca crassidens*, hereafter FKW), Rissо’s dolphin (*Grampus griseus*, hereafter RD), and harbor porpoise (*Phocoena phocoena*, hereafter HP). Morphometric inputs are obtained from high-fidelity three-dimensional CAD reconstructions, and passive drag coefficients are derived from WMLES at three reference speeds. We demonstrate the algebraic origin of body-form invariance in speed retention and mass-specific savings, and show that the *T*_opt_ and *T*_zero_ framework provides a species-independent energetic benchmark for interpreting tag-derived glide durations across cetacean species.

## 2. Results

### 2.1 Optimal and maximum glide durations

*T*_opt_ scales strongly with body size and decreases monotonically with increasing swimming speed for all three species (Table 1; model derivation in §5.3–5.4). At 1.5 m s−1, the FKW optimal glide duration (23.27 s) is 1.73 times that of the RD (13.44 s) and 3.79 times that of the HP (6.14 s), consistent with *T*_opt_ being proportional to *M*_v_ / (*S* · *C*_d_) at fixed speed (§5.5). Active swimming power (*P*_ss_) rises steeply with speed for all species due to the cubic dependence of drag-based propulsive power on velocity. At 1.5 m s−1, *P*_ss_ for the HP (40.9 W) is 18.6% of that for the FKW (219.7 W), but on a mass-specific basis the HP requires 3.79 times more power per kilogram. This identical ratio reflects the same *S* · *C*_d_ /*M*_v_ scaling that governs *T*_opt_, as both quantities depend on the same morphometric group. Glide profitability curves for all three species and speeds are shown in Figure 1. The speed decay profiles underlying these timescales are shown in Figure 2.

**Figure 1.**
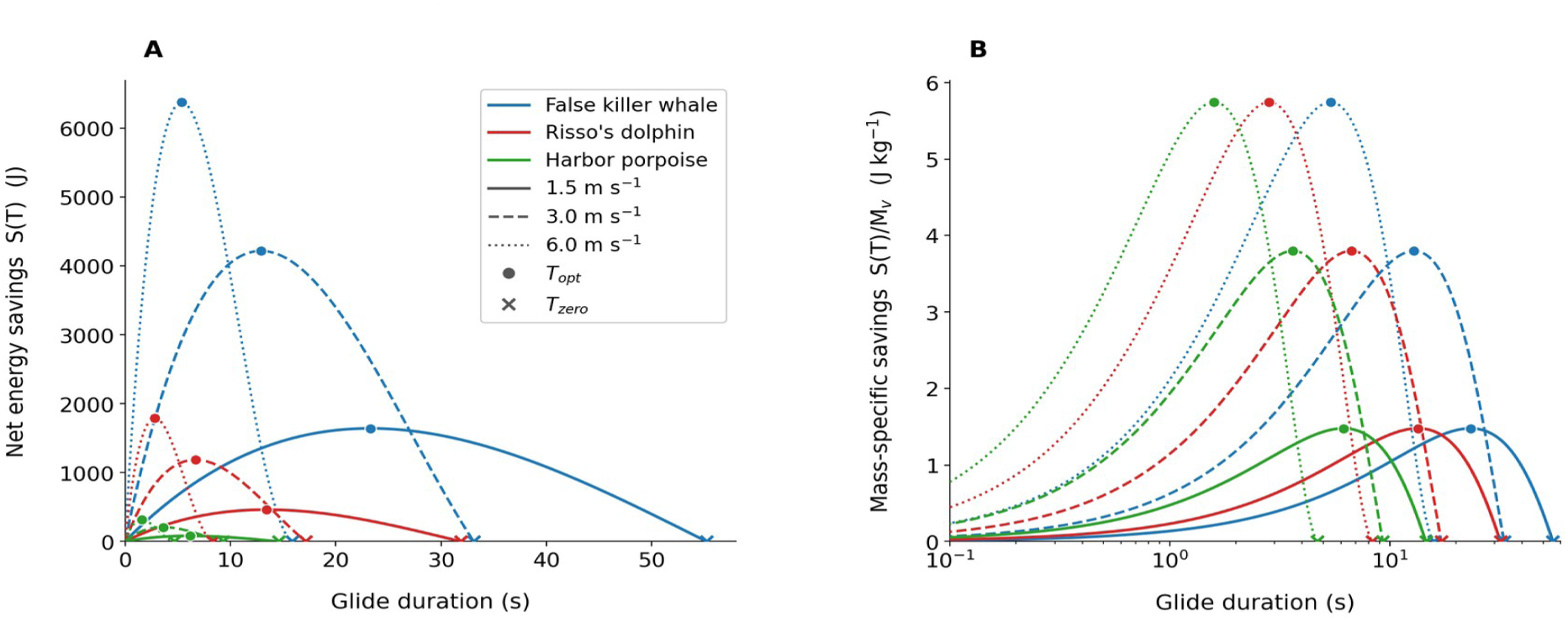
Glide profitability curves for three odontocete species at three swimming speeds (solid, dashed, dotted lines). Filled circles mark *S*(*T*_opt_) = *E*_peak_; crosses mark *S*(*T*_zero_) = 0. **(A)** Absolute net energy savings *S*(*T*) (J), linear time axis. **(B)** Mass-specific savings on a logarithmic time axis. False killer whale, blue; Risso’s dolphin, red; harbor porpoise, green.

**Figure 2.**
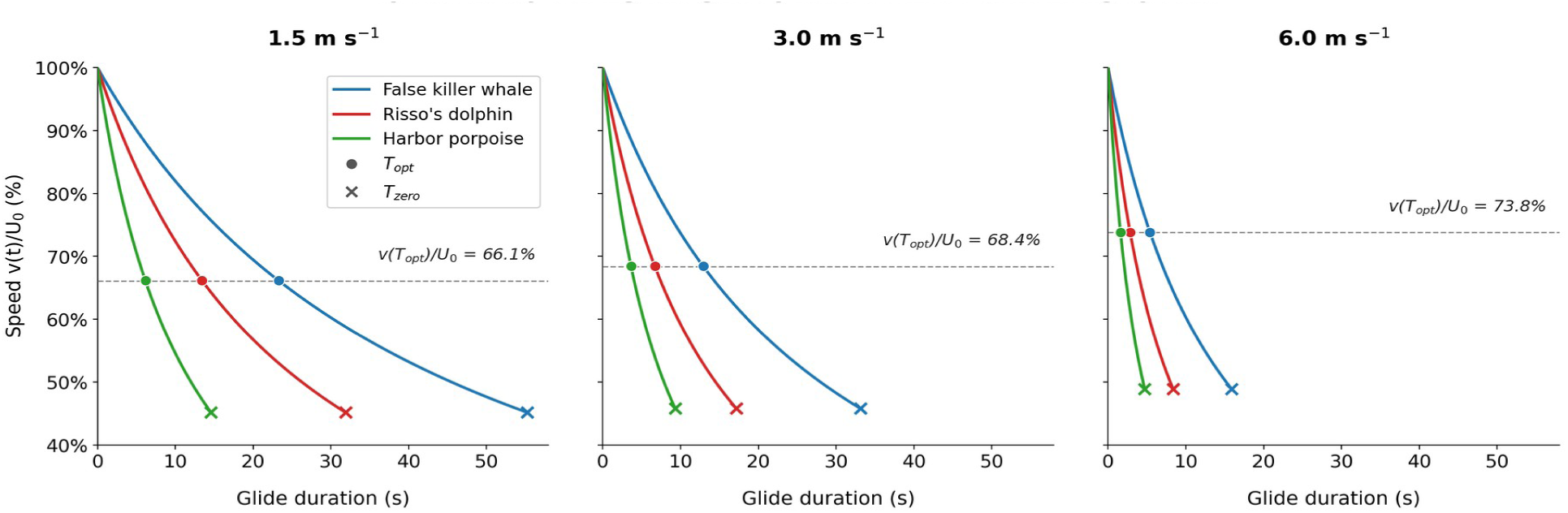
Speed *v*(*t*)/*U*_0_ during the passive glide phase at three swimming speeds. Filled circles mark *T*_opt_; crosses mark *T*_zero_. The dashed line shows the invariant retention level, identical across species at each speed (§2.2).

**Table 1.**
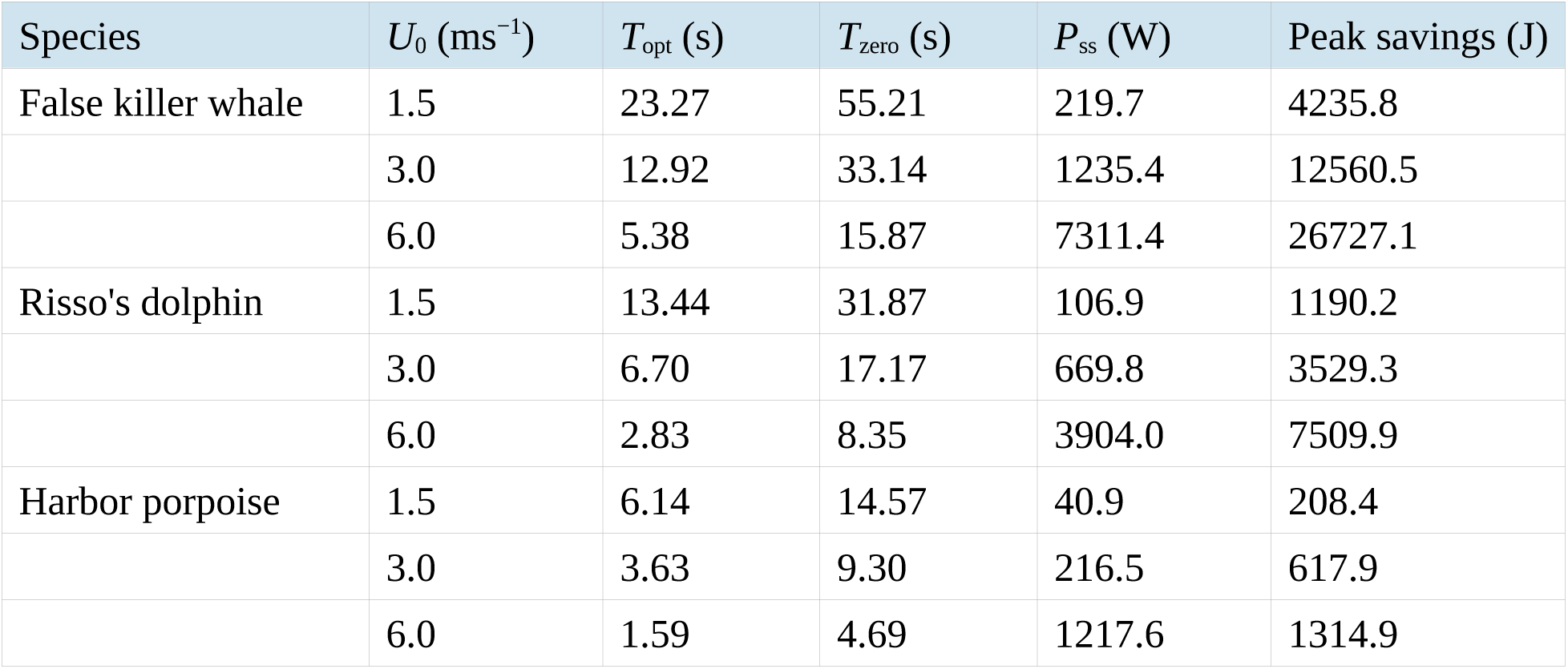
Benchmark values of optimal glide duration (T_opt_), maximum glide duration (T_zero_), active swimming power (P_ss_), and peak energy savings at three reference speeds for all three species.

| Species | $U_0$ (ms<sup>-1</sup>) | $T_{\text{opt}}$ (s) | $T_{\text{zero}}$ (s) | $P_{\text{ss}}$ (W) | Peak savings (J) |
| --- | --- | --- | --- | --- | --- |
| False killer whale | 1.5 | 23.27 | 55.21 | 219.7 | 4235.8 |
|  | 3.0 | 12.92 | 33.14 | 1235.4 | 12560.5 |
|  | 6.0 | 5.38 | 15.87 | 7311.4 | 26727.1 |
| Risso's dolphin | 1.5 | 13.44 | 31.87 | 106.9 | 1190.2 |
|  | 3.0 | 6.70 | 17.17 | 669.8 | 3529.3 |
|  | 6.0 | 2.83 | 8.35 | 3904.0 | 7509.9 |
| Harbor porpoise | 1.5 | 6.14 | 14.57 | 40.9 | 208.4 |
|  | 3.0 | 3.63 | 9.30 | 216.5 | 617.9 |
|  | 6.0 | 1.59 | 4.69 | 1217.6 | 1314.9 |

### 2.2 Body-form invariance of speed retention

The algebraic cancellation of the passive drag terms when substituting the optimal glide duration into the speed-decay law produces a body-form-invariant speed retention function that is identical for all three species at any given speed (Figure S1). At 1.5 m s−1, where *g* = 3.2, all three species retain 66.1% of their initial speed at *T*_opt_. Retention rises gradually with swimming speed, reaching 73.8% at 6.0 m s−1 where *g* = 2.0. This increase reflects the speed-dependence of *g*: at higher speeds, *g* is lower, so *T*_opt_ is reached sooner and the animal retains more speed when it resumes stroking. The three species return identical retention values at each speed, confirming the body-form invariance result (equation 10), and the full speed-dependent curve is provided in Figure S1.

The ratio *T*_zero_/*T*_opt_ ranges from 2.37 at 1.5 m s−1 to 2.95 at 6.0 m s−1, indicating that the window of positive energetic benefit widens relative to the optimal point as speed increases. Because this ratio depends only on *Ψ*(*U*_0_), it is identical across species at each speed. A retention value of approximately 66% at low speeds corresponds to roughly 44% of initial kinetic energy at *T*_opt_, with the remaining 56% dissipated against hydrodynamic drag. This retention value yields a simple energetic prediction: the energetically optimal time to resume stroking is when the animal has decelerated to approximately 66% of its initial speed at low-to-moderate speeds, independently of body size or form. Whether cetaceans glide to this fraction, or overshoot or undershoot it for behavioral or physiological reasons, is a testable prediction for tag-derived kinematic data. Sensitivity bounds incorporating Re- and *C*_d_-corrections for each species are provided in Figure S1 and discussed in §2.5.

### 2.3 Mass-specific peak savings: a second invariant result

In the mass-specific savings expression *E*_peak_/*M*_v_ = *f*(*x*_opt_) · *U*_0_2 / *η*, all terms involving *M*_v_, *S*, *C*_d_, and *ρ* are absent. The result depends only on *g*(*U*_0_), *η*(*U*_0_), and *U*_0_, and is identical across species when *η* is treated as shared. Near-constant *η* across odontocetes is supported by the approximate invariance of dimensionless fluke geometry, including cross-sectional parameters and moderate variation in aspect ratio, across body sizes and species (Pavlov et al., 2021), and is consistent with empirical Froude efficiency estimates of approximately 0.70 to 0.90 (Fish, 1998; Rohr and Fish, 2004). The allometric relationships among *M*_v_, *S*, and *C*_d_ in cetaceans are consistent with the cancellation of these terms in the mass-specific savings expression, but are not required for it. The same invariance applies to any set of bodies for which *g(U)* and *η* are shared. Panel B of Figure 1 illustrates this invariance.

### 2.4 Body form differentiation: absolute glide duration, glide distance, and energetics

While the two invariants above hold at the proportional level, body form creates large differences in the absolute scale of burst-and-glide locomotion. Distance covered during the glide is:

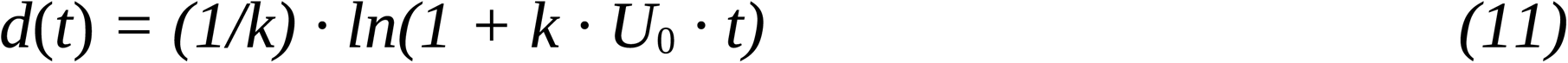

Evaluated at *T*_opt_, glide distance scales with *M*_v_ / (*S* · *C*_d_). At 1.5 m s−1, the FKW covers 28.2 m per optimal glide compared with 7.4 m for the HP, a 3.79-fold difference consistent with the *T*_opt_ ratio at this speed. Energy savings per meter of passive transport are 5.4-fold larger for the FKW (150.3 J m−1) than for the HP (28.0 J m−1), a ratio exceeding the *T*_opt_ ratio because savings per meter scale with *S* · *C*_d_ rather than *M*_v_. The temporal window available for glide optimization also differs sharply. The FKW has approximately 23 s at 1.5 m s−1 to adjust glide timing relative to dive phase or ambient current conditions, whereas the HP has fewer than 7 s before reaching *T*_opt_ and approximately 14.6 s before savings turn negative.

The mechanical cost of transport (COT_mech_) is 3.79 times higher for the HP than for the FKW at 1.5 m s−1 (0.500 versus 0.132 J kg−1 m−1), driven by the higher passive *C*_d_ of the smaller animal. The mass-specific peak savings of 3.82 J kg−1 at 1.5 m s−1 represent 83% of the mass-specific active swimming expenditure over the glide duration for all three species equally, a consequence of the species-invariant efficiency factor *η*_eff_ at this speed (defined in §5.4 and Appendix, equation A3).

### 2.5 Sensitivity of speed retention and mass-specific savings to interspecific *g* variation

The body-form invariance result is conditional on g(U) being approximately species-independent at each absolute speed, a first-order simplification justified by WMLES boundary-layer results showing predominantly turbulent flow at all three benchmark speeds for all three species (§5.3). The Reynolds number spans nearly threefold across the study species at 1.5 m s⁻¹, from HP (Re ≈ 2.3 × 10⁶) through RD (Re ≈ 4.2 × 10⁶) to FKW (Re ≈ 6.6 × 10⁶), and its quantitative effect on speed retention requires direct assessment. Direct measurements of *g* for the study species are not yet available. Published empirical values span a wide range (*g* = 2.7 to 10.6; Webb, 1975; Fish et al., 2014), and values approaching 10 are associated with low-speed conditions in pilot whales (Webb, 1975), where a partially laminar passive boundary layer undergoes near-complete transition under active fluking. This regime is mechanistically distinct from the sustained cruising speeds modeled here, as confirmed by the WMLES results (§5.2).

We assessed sensitivity of speed retention to interspecific *g* variation using two complementary corrections. First, species-adjusted values were computed as *g*_sp_ = *g*_shared_ × (Re_ref_ / Re_sp_)*k*(*U*), where Re_ref_ is the Reynolds number of the bottlenose dolphin calibration reference (*L*_ref_ ≈ 2.5 m; Fish, 1993) and Resp is the species Reynolds number at the same absolute speed. The exponents *k*(*U*) = 0.385, 0.160, and 0.109 at 1.5, 3.0, and 6.0 m s−1 are derived from the Re-dependence of skin-friction scaling in turbulent boundary layers (Schlichting, 1979) and decrease with speed, reflecting the reduced sensitivity of boundary-layer state to Re in the fully turbulent regime. FKW and RD, larger than the calibration reference, receive lower Re-adjusted *g* values, placing their retention above the invariant curve by 6.26 percentage points for FKW and 1.63 percentage points for RD at 1.5 m s−1. HP receives higher Re-adjusted *g*, placing its retention below the invariant curve (HP: −4.3 percentage points at 1.5 m s−1). All deviations converge to within 1 to 2 percentage points of the invariant curve at 6.0 m s−1. Second, each species’ *g* was scaled by the inverse ratio of passive drag coefficients relative to FKW: *g*_sp_ = *g*shared × (*C*_d_,FKW / *C*_d_,species). This correction assumes that *C*_d_,active is approximately species-independent at a given absolute speed. Physically, this would hold if the fluke-stroke kinematics impose a sufficiently strong and uniform turbulence perturbation on the boundary layer that any residual species-specific laminar fraction is completely disrupted regardless of Re. Under this assumption, *g* = *C*_d_,active /*C*_d_,passive scales inversely with the passive drag coefficient. The alternative limit, in which *C*_d_,active and *C*_d_,passive scale proportionally across species, would leave *g* species-independent. The two corrections together bracket the plausible range. Under the inverse-scaling assumption, this yields *C*_d_-adjusted *g* for RD of approximately 3.00, 2.27, and 1.71 and for HP of approximately 2.29, 2.06, and 1.61 at 1.5, 3.0, and 6.0 m s−1, all below the shared model values and placing speed retention above the invariant curve for both species.

For HP, the Re-correction yields 61.9% at 1.5 m s−1 (−4.3 percentage points) while the *C*_d_-correction yields 75.0% (+8.8 percentage points). Because the two corrections act in opposing directions for HP, the invariant curve lies within the HP envelope. The envelope width is approximately 13 percentage points at 1.5 m s−1, narrowing to approximately 9 percentage points at 6.0 m s−1. For FKW and RD, both corrections act in the same direction, placing the invariant curve below their plausible retention range. For RD, whose body length is close to the calibration reference (2.5 m), both corrections are small and nearly equal at 1.5 m s⁻¹. The Re-correction weakens faster with increasing speed, governed by the declining exponent *k(U)*, so the two RD bounds cross near 1.5 m s⁻¹ and diverge thereafter, as shown in Figure S1.

A second departure from exact invariance arises within each glide. Because passive *C*_d_ rises as the animal slows, holding it at the initial-speed value overstates retention by up to 2.4 percentage points and introduces a cross-species spread of up to 1.6 percentage points, both largest near 3.0 m s−1 where the *C*_d_(*U*) curve is steepest. These bounds are smaller than the *g*-driven envelope above and share its Reynolds-number origin.

A third mechanism driving *g* variation is tail-beat frequency (TBF). Higher TBF elevates *g* by destabilizing the boundary layer more frequently, independently of Reynolds number or passive *C*_d_. TBF scales inversely with body length across cetaceans (Sato et al., 2007; Gough et al., 2019), so HP exhibits higher TBF than FKW at any given absolute speed, acting cumulatively with the Reynolds number correction. Neither mechanism affects the algebraic cancellation established in §3.1 and §5.5, which holds for any *g*(*U*) including species-specific values. The Re-correction and TBF scaling act in the same direction, both increasing g for HP relative to the shared baseline. The *C*_d_-correction acts in the opposite direction. The net effect on HP’s retention depends on which mechanism dominates, a question that cannot be resolved without direct measurement of active drag. For FKW and RD, all three mechanisms act in the same direction: lower TBF, higher Re, and lower *C*_d_,passive relative to HP all reduce *g* below the shared baseline, placing their retention consistently above the invariant curve. Measurements of active drag on each study species remain the high-priority experimental target for resolving the net direction of these competing effects in HP and confirming the systematic upper-bound interpretation for FKW and RD.

The same *g* variation propagates more strongly to the mass-specific savings than to speed retention. Speed retention depends on *g* through the factor 1/(1 + *x*_opt_), which damps the response, whereas the savings depend on *g* through *f*(*x*_opt_), in which *g* enters directly as a multiplier. Evaluated at the benchmark speeds, the logarithmic sensitivity of the savings to *g* is six to eight times that of retention. Applying the same Re- and *C*_d_ -corrections used above for HP, the mass-specific savings range from 50 percent above the shared-*g* value to 60 percent below it at 1.5 m s−1. At 6.0 m s−1 the range is 18 percent above to 58 percent below. The cross-species equality of the savings is thus a weaker numerical claim than that of retention, holding within tens of percent rather than the nine to thirteen percentage points of the retention envelope reported above. The algebraic cancellation of *M*v, *S*, *C*_d_, and *ρ* in the mass-specific savings expression (§2.3, §5.4) is unaffected, exactly as for retention, since it holds for any *g*(*U*). Direct measurement of active drag, already identified above as the priority for resolving the retention bounds, would equally constrain the savings.

## 3. Discussion

Locomotor cost in swimming vertebrates scales steeply with body size (Schmidt-Nielsen, 1972; Watanabe et al., 2011; Williams, 2022; Glarou et al., 2025), and the glide timescales reported here behave accordingly, with *T*_opt_ differing by nearly fourfold across the species studied through the body-size dependence of the deceleration timescale 1/*α*. Two quantities derived from the glide, however, do not scale with body form. Speed retained at *T*_opt_ is identical across species at any given speed, and mass-specific peak energy savings are independent of body morphometry. The invariance follows from a single dimensionless group, *x*_opt_, which absorbs the morphometric and hydrodynamic parameter space of the glide phase.

This invariance arises because the cancellation of morphometric and hydrodynamic terms holds unconditionally for any *g*(*U*), regardless of whether *g* is species-specific. The equality of speed retention values across species holds when *g*(*U*) is treated as approximately species-independent at a given absolute speed, a simplification assessed quantitatively in §2.5. The energetic advantage of gliding extends across a wide size range because *g* depends primarily on boundary-layer state, governed by Reynolds number rather than body size alone (Borazjani and Sotiropoulos, 2008). Body size and form set the absolute scale of burst-and-glide locomotion, including *T*_opt_, glide distance, and energy saved per cycle, but not the mass-specific energetics of either phase. The active and passive phases are energetically coupled through propulsive efficiency *η*, which is constrained by the Strouhal number (Rohr and Fish, 2004; Saadat et al., 2017), and through *η*_eff_, the species-invariant factor that governs the fraction of active-phase savings recovered during each glide (§5.4).

*x*_opt_ exemplifies the Buckingham π theorem (Buckingham, 1914), collapsing the dimensional parameter space of body morphology and hydrodynamic parameters into a single dimensionless group that fully determines the optimal glide duration independently of species. The underlying principle of invariance under a change of units was extended by Liu et al. (2025b) through symmetry transformations of the governing partial differential equations. The authors identified Re, Fr, St, and a newly recognized elastic analogue Em as the dimensionless groups governing locomotion dynamics across the full diversity of animal life, from microorganisms to large mammals. *x*_opt_ therefore complements this family of locomotion-governing dimensionless groups by defining the energetic optimum of the glide phase.

Intermittent locomotion, alternating active propulsion with gliding, is a convergent strategy across aquatic and aerial vertebrates spanning many orders of magnitude in body mass, reflecting physical and physiological principles that broadly favor energetically efficient transport (Weihs, 1974; Rayner, 1985; Williams et al., 2000; Gleiss et al., 2011; Liu et al., 2025a). Prior work has identified four size-independent optima for the active phase of cetacean locomotion. The Strouhal number of undulatory propulsion is constrained to a narrow, efficiency-maximizing range regardless of body size (Saadat et al., 2017; Rohr and Fish, 2004). The normalized energy cost of active swimming is approximately constant across swimmers and fliers spanning 10–20 orders of magnitude in mass (Bale et al., 2014). The kinematics of inertial undulatory swimmers from millimeter-scale larvae to 30-metre whales follow a universal scaling relation (Gazzola et al., 2014). The COT-minimizing swim speed is nearly size-independent across marine vertebrates (Watanabe et al., 2011). These optima all characterize the active phase, whereas the passive phase lacked an equivalent description. The present results fill this gap, identifying speed retention at *T*_opt_ and mass-specific peak energy savings as the passive-phase analogues of the active-phase optima. The mass-specific energetics of both phases are governed by hydrodynamic dimensionless groups (*g*, St, and *x*_opt_), suggesting dynamic similarity of the full burst-and-glide cycle across the odontocete size range at any given swimming speed. This coherence is consistent with a convergent evolutionary trajectory toward COT minimization across a wide size range, arising from hydrodynamic principles rather than from size-specific morphological tuning (Gleiss et al., 2011).

These invariants enable direct cross-species comparison of glide duration data collected from bio-logging tags. The invariant results identify the energetic optimum within the trade-off between momentum loss and energy savings during gliding. Once *T*_opt_ is computed for each species from its morphometric and drag parameters, tag-derived glide durations expressed as *T*/*T*_opt_ are directly comparable across species regardless of body form. Expressing them this way does not eliminate the need for species-specific morphometric inputs, since *T*_opt_ scales as *M*_v_/(*S*·*C*_d_) and must be computed individually, but the normalization ensures that deviations from *T*/*T*_opt_ = 1 or *T*/*T*_zero_ = 1 reflect behavioral or environmental context rather than differences in body form, establishing a common energetic scale for cross-species comparison. Such deviations may arise from buoyancy changes during diving (Skrovan et al., 1999; Miller et al., 2004), prey pursuit (Arranz et al., 2019), social interaction, acoustic constraints, or responses to anthropogenic disturbance (Wisniewska et al., 2018). Because the model depends only on quadratic passive drag, it extends in principle to any odontocete for which passive drag coefficients and an estimate of *g*(*U*) are available, placing cross-species comparisons of gliding behavior on a common energetic basis. The same invariants constrain the burst-and-glide duty cycle, DC = *T*_active_ / (*T*_active_ + *T*_opt_), and Floryan et al. (2017) showed that the COT-minimizing duty cycle depends on the drag ratio and swimming speed, a finding directionally consistent with the present framework.

The framework rests on several simplifying assumptions. Gliding is treated as occurring in still water without buoyancy or gravitational assistance, so descent-assisted gliding in deep divers such as the FKW may constitute a qualitatively different regime from the horizontal burst-and-glide modeled here (Skrovan et al., 1999; Miller et al., 2004; Arranz et al., 2019). Hydrodynamics are treated as quasi-steady, with drag coefficients from steady gliding simulations applied throughout deceleration, thereby neglecting acceleration-dependent added-mass forces and transient boundary-layer effects at fluke-stroke termination. The coefficients of *Φ*(*U*_0_) and *Ψ*(*U*_0_) are treated as species-independent and anchored to *Tursiops truncatus* data from Fish (1993), so direct validation against high-resolution tag data for each species would strengthen confidence in their generality. Sensitivity to the *g*(*U*) assumption, the key limitation, is quantified in §2.5, and species-specific variation in *g* shifts the absolute values of *T*_opt_ and *T*_zero_ without affecting the algebraic cancellation that underlies the two invariant results. The model does not represent maneuvering, acoustic, or thermoregulatory loads on nominally passive glides, and the COT-minimization objective applies to transit swimming rather than to prey pursuit, station-keeping, or social interaction.

Despite these limitations, the two passive-phase optima complement the established active-phase invariants, contributing to a unified, scale-independent description of the full burst-and-glide cycle. The principal remaining uncertainty is the species-dependence of *g*, and the CAD-CFD approach applied here to passive gliding is the natural next step toward resolving it for active swimming, given the obvious experimental and logistical difficulties in studies of free-swimming species. A follow-up study applies this framework to bio-logging tag data from the same species to estimate departures from energetically optimal *T*_opt_ and *T*_zero_ associated with behavioral patterns.

## 4. Conclusions

We developed a glide-phase energetics model for three odontocete cetaceans spanning a 20-fold mass range. Two invariant results emerge from the analysis. First, speed retention at the energetically optimal glide duration (*T*_opt_) is analytically independent of *M*_v_, *S*, *C*_d_, and *ρ*, depending only on *g*(*U*) and swimming speed. Second, mass-specific peak energy savings at *T*_opt_ are invariant across species, arising from the same algebraic cancellation of morphometric and hydrodynamic parameters under the assumption that propulsive efficiency *η* is shared, a condition supported by the approximate invariance of dimensionless fluke geometry across body sizes and species (Pavlov et al., 2021).

Body form differentiates burst-and-glide locomotion by absolute glide duration and glide distance, both up to 3.79-fold between species. Both invariants arise from the proportionality of *T*_opt_ to the passive deceleration timescale 1/*α*, which causes all terms comprising the passive drag constant *k* to cancel algebraically. Sensitivity analysis incorporating Re- and *C*_d_-corrections shows that two sources of interspecific *g* variation act in opposing directions for the smallest species, with the invariant curve falling within a band of approximately 13 percentage points at low speeds.

The present results complement prior size-independent optima for the active phase of cetacean locomotion, including near-size-invariant COT-minimizing speeds (Watanabe et al., 2011), universal Strouhal number bounds (Saadat et al., 2017; Rohr and Fish, 2004), universal kinematics scaling across inertial undulatory swimmers (Gazzola et al., 2014), and approximate constancy of normalized locomotor energy cost (Bale et al., 2014). The glide-phase energetics framework developed here extends to any cetacean species for which morphometric and drag data are available, yielding species-specific *T*_opt_ and *T*_zero_ as metrics of gliding efficiency on a common energetic scale. Together, these active- and passive-phase invariants form a scale-independent energetic framework for burst-and-glide locomotion, consistent with convergent evolution toward COT minimization across a wide size range.

## 5. Materials and Methods

### 5.1 CAD modeling and morphometry

Three odontocete species were selected to represent a broad range of body sizes within the suborder: FKW, RD, and HP. These species span approximately 20-fold variation in body mass (54.6 to 1109.9 kg) and 2.9-fold variation in body length (1.59 to 4.65 m), providing a wide dynamic range for detecting body-form effects on glide energetics.

Body measurements of stranded animals obtained from stranding networks and the literature were used to create realistic CAD models. Body condition was consistent with Decomposition Condition Code 2 (freshly dead). Adult females were selected based on body length, with a standard deviation of less than 4%. Outliers in body mass and girth, including pregnant and exhausted individuals, were excluded. CAD models were constructed using SolidWorks® software at a 1:1 scale. Fin planes were oriented orthogonally to the frontal YZ plane to achieve zero angle of attack with respect to the flow direction. The dihedral angle ranged from 153° to 171°, and fin sweep angles were adjusted where necessary to achieve near-zero-lift conditions. Photos and videos of free-swimming individuals were used to select a straightened body posture characteristic of the gliding phase. The resulting morphometric parameters are summarized in Table 2. Virtual mass *M*_v_ = *M*_b_ + *M*_added_, where *M*_b_ is body mass and *M*_added_ is the added mass of water accelerated with the body during axial motion. For a neutrally buoyant body gliding along its long axis, axial added mass is given by potential flow theory for a prolate spheroid as *M*_added_ = *C*_a_ · *ρ* · *V*, where *V* is displaced volume and *C*_a_ decreases strongly with fineness ratio FR = *L*/*D* (Lamb, 1932; Brennen, 1982). Fineness ratios measured directly from the CAD models are FR = 6.04 (FKW), 5.48 (RD), and 5.09 (HP), yielding *C*_a_ = 0.045, 0.055, and 0.062 respectively from potential flow theory, corresponding to *M*_added_ ≈ 4.5–6.2% of body mass for a neutrally buoyant animal. A shared value of 5% was adopted as a representative central estimate consistent with this range, giving *M*_v_ = 1.05 · *M*_b_ (FKW: *M*_b_ = 1057.0 kg, *M*_v_ = 1109.9 kg; RD: *M*_b_ = 297.0 kg, *M*_v_ = 311.9 kg; HP: *M*_b_ = 52.0 kg, *M*_v_ = 54.6 kg). The commonly cited value of approximately 20% applies to transverse rather than axial motion and is therefore not appropriate here (Gero, 1952; Webb, 1975).

**Table 2.** Species-specific morphometric and hydrodynamic parameters used in the energetics model.

| Parameter | Unit | FKW | RD | HP |
| --- | --- | --- | --- | --- |
| Body length | m | 4.65 | 2.95 | 1.59 |
| Virtual mass (Mv) | kg | 1109.85 | 311.85 | 54.6 |
| Wetted surface area (S) | m<sup>2</sup> | 8.06 | 3.68 | 1.076 |
| Cd at 1.5 m s<sup>-1</sup> | — | 0.00394 | 0.0042 | 0.0055 |
| Cd at 3.0 m s<sup>-1</sup> | — | 0.0032 | 0.0038 | 0.0042 |
| Cd at 6.0 m s<sup>-1</sup> | — | 0.00295 | 0.00345 | 0.00368 |
| g at 1.5 m s<sup>-1</sup> | — | 3.2 | 3.2 | 3.2 |
| g at 3.0 m s<sup>-1</sup> | — | 2.7 | 2.7 | 2.7 |
| g at 6.0 m s<sup>-1</sup> | — | 2.0 | 2.0 | 2.0 |
| $\eta$ at 1.5 m s<sup>-1</sup> | — | 0.80 | 0.80 | 0.80 |
| $\eta$ at 3.0 m s<sup>-1</sup> | — | 0.78 | 0.78 | 0.78 |
| $\eta$ at 6.0 m s<sup>-1</sup> | — | 0.72 | 0.72 | 0.72 |
| Water density ($\rho$) | kg m<sup>-3</sup> | 1000 | 1000 | 1000 |

### 5.2 Passive drag coefficients

Passive drag coefficients (*C*_d_) at three reference speeds (1.5, 3.0, and 6.0 m s⁻¹) were obtained from wall-modeled large eddy simulations (WMLES). Simulations were performed using charLES (Brès et al., 2018), which solves the compressible Navier–Stokes equations in the low-Mach regime using an explicit, unstructured, finite-volume scheme. An isentropic (Helmholtz) approximation retains a bounded but finite acoustic speed, so that the time step is governed by the convective rather than the acoustic CFL constraint. At the freestream speeds considered, the Mach number *M* = *U*_∞_/*c*, with *c* ≈ 1481 ms^−1^, does not exceed 0.004. Compressibility corrections scale as *M*² and are therefore below 2 × 10⁻⁵, so the low-Mach solution reproduces the incompressible drag coefficient to within this bound. The solver is second-order accurate in space and third-order accurate in time. All simulations were performed using the GPU-accelerated parallel executable.

The unresolved turbulent scales are modeled using the Vreman subgrid-scale model (Vreman, 2004). An algebraic wall model was employed to provide the wall shear stress closure. A first-point matching approach was adopted throughout, and this formulation has been shown to avoid log-layer mismatch (Kawai et al., 2012). The first off-wall matching point corresponds to y⁺ in the range 12–147 across all configurations, with the local value varying along the body length and increasing with freestream speed (Table 3); these values lie within the range typically adopted in wall-modeled LES, for which the algebraic wall model provides the wall-stress closure.

**Table 3.**
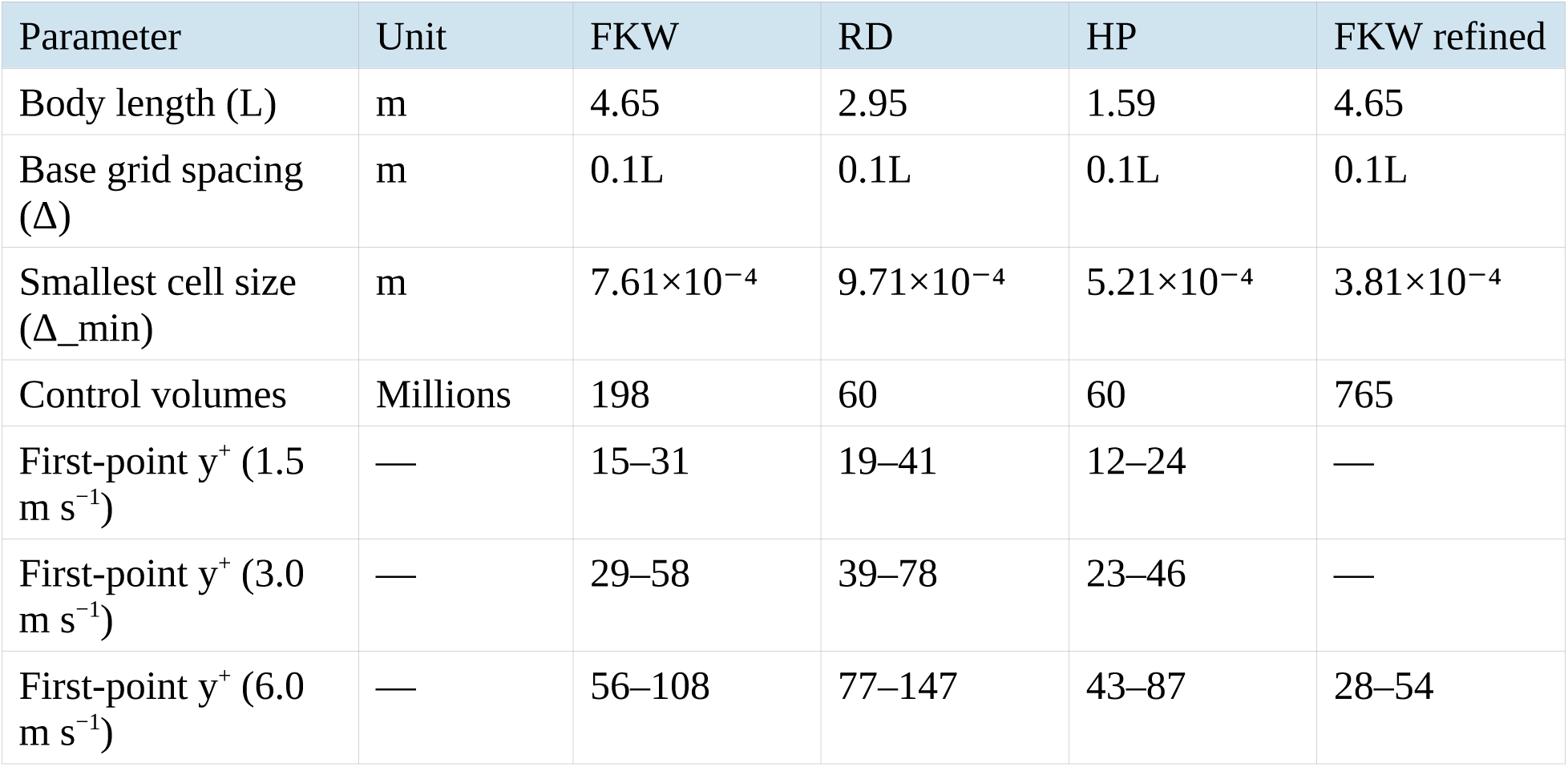
Grid characteristics for each configuration: body length, base spacing, smallest cell size, control volume count, and first-point y⁺ ranges at the three benchmark speeds.

The computational grids are based on Voronoi diagrams generated from a hexagonally close-packed (HCP) lattice. Local grid refinement is achieved through isotropic subdivision by factors of two, controlled via refinement levels. Near-wall refinement is imposed through layered refinement zones surrounding the body surface. For all configurations, the computational domain extends 5*L* upstream, 10*L* downstream, and 5*L* in both lateral and vertical directions, where *L* is the body length. The base grid spacing is defined as Δ = 0.1*L*, with progressive refinement toward the body using up to 10 near-wall layers.

At the inlet, a uniform freestream velocity *U*∞ is prescribed. The lateral and vertical boundaries are treated as slip walls to mimic far-field conditions. At the outlet, a non-reflecting boundary condition is imposed with a reference pressure set to zero. The characteristic length scale used in the outlet formulation is taken as the body length, *L*_ref_ = *L*, ensuring consistency with the physical scale of the problem and avoiding artificial pressure gradients across the domain.

The main characteristics of the computational grids for each configuration are summarized in Table 3. The FKW, Risso’s dolphin, and harbor porpoise cases employ maximum refinement levels of 9, 8, and 8, respectively, resulting in grids of approximately 198, 60, and 60 million control volumes. The corresponding minimum cell sizes are 7.61 × 10⁻⁴ m, 9.71 × 10⁻⁴ m, and 5.21 × 10⁻⁴ m, respectively. Grid refinement is concentrated in the vicinity of the body and progressively relaxed toward the far field through layered isotropic refinement.

The simulations were advanced until a statistically stationary state was reached. The drag coefficient was computed by averaging over the final 20 flow-through times, *T*_ft_ = *L*/*U*∞, using instantaneous force data from the solver.

An additional higher-resolution simulation was carried out for the FKW configuration at *U*∞ = 6 m s⁻¹ to quantify the uncertainty in the predicted drag coefficient due to mesh resolution. This case was selected because it corresponds to the highest Reynolds number considered and is therefore expected to require the finest resolution, providing a conservative bound for all other cases. This refined mesh employed one additional refinement level (level 10), resulting in approximately 765 million control volumes and a minimum cell size of 3.81 × 10⁻⁴ m (see Table 3). The difference in *C*_d_ between the baseline and refined simulations was found to be approximately 1%, indicating low sensitivity to grid resolution. This variation was therefore used as a conservative estimate of the mesh-induced uncertainty and applied to all cases.

The drag coefficients at the three benchmark speeds (Table 2) were interpolated to intermediate velocities using piecewise cubic Hermite interpolating polynomials (PCHIP) in 0.1 m s−1 steps across 0.5 to 6.0 m s−1. PCHIP preserves monotonicity and avoids spurious oscillations, appropriate for the smooth, monotonically decreasing *C*_d_(*U*) relationship expected from Re-dependent skin-friction scaling in turbulent boundary layers (Schlichting, 1979).

### 5.3 Active drag factor and propulsive efficiency

The active drag factor *g*, introduced by Weihs (1974) and operationalized empirically by Fish (1993) and Fish et al. (2014), governs the ratio of fluking drag to passive gliding drag at the same speed. We adopt the formulation of Weihs (1974) and Floryan et al. (2017), assigning *g* shared baseline values across species as a first-order simplification. The specific baseline values (*g* = 3.2, 2.7, and 2.0 at 1.5, 3.0, and 6.0 m s⁻¹) are calibrated to the empirical value of *g* = 3.2 reported for *Tursiops truncatus* by Fish (1993), with the speed dependence derived from boundary-layer scaling arguments (Schlichting, 1979). The treatment of *g* as approximately species-independent is a simplifying assumption of this paper whose consequences are assessed in §2.5. Propulsive efficiency *η* values are drawn from the empirical ranges reported by Fish (1998) and Rohr and Fish (2004) and applied here without species-specific fitting.

The magnitude of *g* is governed by the extent to which active fluking disrupts the boundary layer relative to its state during passive gliding (Fish and Lauder, 2006; Fish et al., 2014). At lower speeds and for smaller animals, the Reynolds number is reduced and a larger fraction of the wetted surface may remain laminar during passive gliding, so that active fluke strokes destabilize the flow over a greater proportion of the body. As speed increases, the passive boundary layer transitions toward turbulence, reducing the marginal turbulence penalty of active fluking and decreasing *g*. The simulations indicate predominantly turbulent boundary layers at the lowest benchmark speed (1.5 m s−1) for all three species, with a residual laminar fraction that is smaller for the larger-bodied FKW (Figure 3). This residual laminar fraction is expected to make *g* slightly higher for HP than for FKW at a given absolute speed, since active fluking disrupts a proportionally larger laminar region (Fish and Lauder, 2006). Treating *g* as approximately species-independent therefore acknowledges but does not eliminate this interspecific variation. Propulsive efficiency *η* was assigned values of 0.80, 0.78, and 0.72 at 1.5, 3.0, and 6.0 m s−1, reflecting the monotonic decline in Froude efficiency with increasing reduced frequency across odontocetes (Fish, 1998; Rohr and Fish, 2004). These values lie within the empirical range of approximately 0.70 to 0.90 reported across odontocetes and represent the declining trend across the modeled speed range. No species-specific fitting was performed. The body-form invariance of mass-specific savings (§2.3) holds for any *η* that is treated as shared across species, independently of the specific values chosen. Both *g* and *η* were interpolated across the full velocity range using PCHIP.

**Figure 3.**
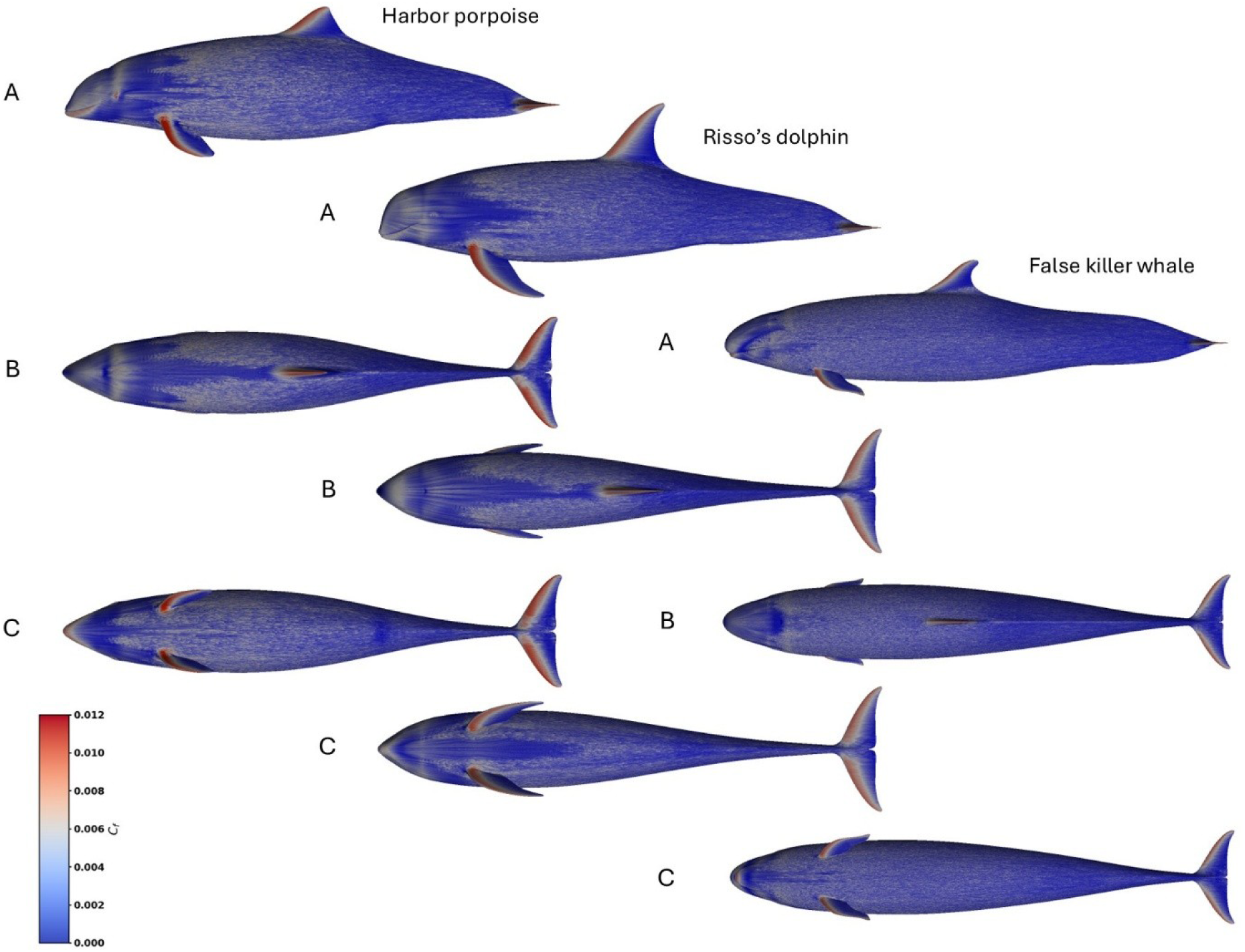
Wall-modeled large eddy simulation (WMLES) at *U*_0_ = 3 m s^−1^. CAD models of harbor porpoise *Phocoena phocoena*, Risso’s dolphin *Grampus griseus*, and false killer whale *Pseudorca crassidens* are scaled to the same size. Color contours show skin-friction coefficient *C*_f_. A – lateral view, B – dorsal view, and C – ventral view.

### 5.4 Glide energetics model

The energetics framework follows the quadratic-drag burst-glide formulation of Weihs (1974), extended here in three respects. First, drag coefficients and propulsive efficiency are treated as velocity-dependent across speeds rather than constant. Second, the optimal glide duration is expressed through an explicit velocity-dependent correction factor Φ(*U*_0_) derived and calibrated in this paper. Third, a closed-form expression for the maximum glide duration *T*_zero_ is introduced here for the first time.

#### Mass-drag constant

The passive deceleration profile is characterized by the mass-drag constant:

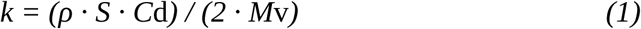

where *ρ* is seawater density (1025 kg m−3), *S* is wetted surface area (m2), *C*_d_ is the passive drag coefficient, and *M*_v_ is virtual mass (the sum of body mass and added mass of entrained water) as defined and tabulated in §5.1 (Table 2). The constant *k* has units of m−1 and captures how rapidly a gliding animal decelerates per unit distance travelled (denoted α in Weihs, 1974).

#### Deceleration rate

The initial deceleration rate at speed *U*_0_ is:

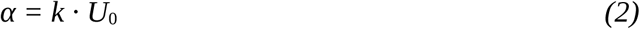

with units of s−1. This is the instantaneous rate of fractional speed loss at the onset of the glide.

#### Optimal glide duration

The glide duration that maximizes net energy savings is:

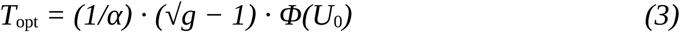

The factor *(√g − 1)* emerges analytically from the Weihs (1974) framework as the primary driver of the optimum, and equals zero when *g* = 1, correctly predicting no glide benefit when active and passive drag are equal. The correction factor Φ(*U*_0_) = 0.58 + 0.046 *U*_0_ is derived in this paper from a first-order Taylor expansion of the full optimization condition (Appendix) and calibrated to *Tursiops truncatus* data from Fish (1993). No additional species-specific fitting was performed. The constant 0.58 is dimensionless and 0.046 has units s m−1. The derivation of *Φ*(*U*_0_) and *Ψ*(*U*_0_) is provided in the Appendix.

#### Maximum glide duration

The glide duration beyond which cumulative savings become negative is:

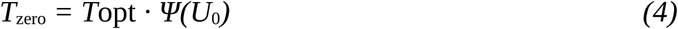

Here Ψ(*U*_0_) = 2.18 + 0.128 · *U*_0_ is a linear function fitted to numerically computed zero-savings values at the three benchmark speeds (Appendix). This closed-form expression is introduced in this paper. The constant 2.18 is dimensionless and 0.128 has units s m−1. Since Ψ exceeds 2.18 for all modeled speeds, an animal can always glide for more than double *T*_opt_ before entering energetic deficit.

#### Steady swimming power and cost of transport

Active swimming power at speed *U* is:

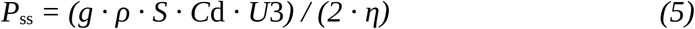

Mass-specific mechanical cost of transport is COT_mech_ = *P*_ss_ / (*M*_v_ · *U*), in J kg−1 m−1. This quantity represents the hydrodynamic propulsion cost normalized per unit mass and distance, and does not include basal metabolic rate.

#### Peak energy savings

The maximum net energy saved during an optimal glide is derived from the savings function *f*(*x*opt) defined in the Appendix (equation A3). In dimensional form:

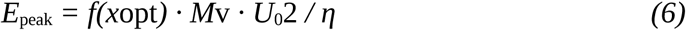

where *f*(*x*_opt_) = *g* · *x*_opt_ − ½(1 − 1/(1 + *x*_opt_)2) and *x*_opt_ = (√*g* − 1) · *Φ*(*U*_0_) is the species-invariant dimensionless glide parameter (see §5.5). The first term of *f* represents the mechanical energy saved during the glide. The second term represents the mechanical work required to re-accelerate from *v*(*T*_opt_) back to *U*_0_. Equivalently, *E*_peak_ = *P*_ss_ · *T*_opt_ · *η*_eff_, where *η*_eff_ = *f*(*x*_opt_) / (*g* · *x*_opt_) is a speed-dependent, species-invariant factor (0.83 at 1.5 m s−1, declining to 0.68 at 6.0 m s−1) that reflects the fraction of potential active-phase savings recovered after re-acceleration losses. This factor is distinct from the propulsive efficiency *η* (Appendix, equation A3). Panel B of Figure 1 shows that mass-specific savings are identical across species at each speed, reaching 3.82, 11.32, and 24.08 J kg−1 at 1.5, 3.0, and 6.0 m s−1 respectively.

#### Passive speed decay

During a glide the only streamwise force is quadratic drag, so the equation of motion is *M*_v_ d*v*/d*t* = −½ *ρ S C*_d_ *v*². Integrating from the initial speed *U*_0_ gives the hyperbolic decay law:

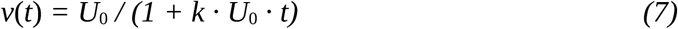

where *k* = *ρ S C*_d_ / (2 *M*_v_) is the passive drag constant (equation 1; Weihs, 1974). Equation (7) is the closed-form solution when *C*_d_ is held constant. Within a single glide *C*_d_ is fixed at its value at *U*_0_. Because *C*_d_ rises as the animal decelerates, holding it constant understates drag late in the glide and slightly overstates speed retention, an effect bounded in §2.5.

### 5.5 Derivation of speed retention at optimal glide duration

The body-form invariance of speed retention is a novel result of this paper. We derive it here by substituting the expression for *T*_opt_ (equation 3) into the speed-decay law (equation 7) and showing that all terms comprising the passive drag constant *k*, including *M*_v_, *S*, *C*_d_, and *ρ*, cancel algebraically. The speed retained at *T*_opt_ is obtained by substituting equation (3) into equation (7), evaluating at *t* = *T*_opt_:

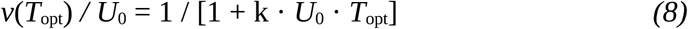

Substituting equations (1) and (2) into (3) gives:

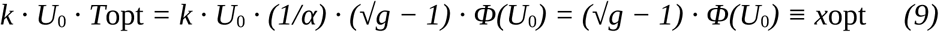

Since *α* = *k* · *U*_0_ (equation 2), the product *k* · *U*_0_ · (1/*α*) = 1 identically, and all terms involving *M*_v_, *S*, *C*_d_, and *ρ* cancel. The dimensionless glide parameter *x*_opt_ depends only on *g*(*U*_0_) and *U*_0_, and is therefore species-invariant. The retention fraction reduces to:

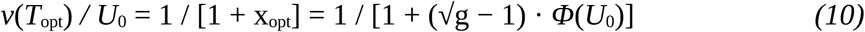

This cancellation follows directly from equations (1), (2), (3), (7), (8), and (9) and holds unconditionally for any *g*(*U*) and any morphometric parameter values. Under the additional assumption that *g*(*U*) is approximately species-independent at a given absolute speed (§5.3), equation (10) yields identical retention fractions for all three species at each swimming speed regardless of body mass, wetted surface area, drag coefficient, or fluid density. The sensitivity of the retention result to interspecific *g* variation is assessed in §2.5.

## Appendix Glide profitability function and structure of *Φ(U*_0_) and *Ψ(U*_0_)

### Glide profitability function

The glide profitability function *S*(*T*) characterizes the net energy saved by gliding passively for duration *T* compared with continuous active swimming over the same time interval at *U*_0_. Three conditions constrain its shape: *S*(0) = 0 (no savings at zero glide duration), *S*(*T*_opt_) = *E*_peak_ (the savings are maximized at the optimal glide duration), and *S*(*T*_zero_) = 0, meaning that at *T*_zero_ the cumulative net advantage of gliding over continuous swimming has been fully extinguished. At this point, the energy cost of re-accelerating from the low speed reached after a long glide equals the energy saved by not fluking, and any further extension of the glide becomes energetically counterproductive. A constrained cubic of the form:

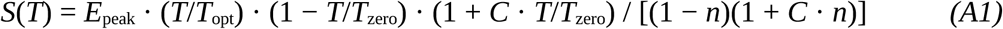

satisfies all three conditions exactly for any *E*_peak_, *T*_opt_, and *T*_zero_. Here *n* = *T*_opt_/*T*_zero_ and the shape parameter:

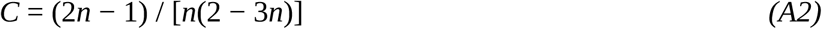

is uniquely determined by requiring d*S*/d*T* = 0 at *T* = *T*_opt_, enforcing the peak at the optimal point regardless of speed or species. Because *n* < 0.5 for all speeds modeled here (the optimal glide duration is always less than half the maximum glide duration), *C* is negative, producing curves that rise steeply to *T*_opt_ and decay more gradually toward *T*_zero_, consistent with the physical expectation that savings accumulate rapidly at the onset of the glide when speed is high but diminish progressively as the animal decelerates. The ratio *n* = *T*_opt_/*T*_zero_ = 1/*Ψ*(*U*_0_) takes values of approximately 0.42, 0.39, and 0.34 at 1.5, 3.0, and 6.0 m s−1 respectively, yielding *C* ≈ −0.51, −0.68, and −0.97 at the same speeds. Figure 1 shows the resulting profitability curves for all three species at all three reference speeds.

### Dimensional peak savings

Substituting *T* = *T*_opt_ into the dimensional form of the net mechanical energy balance, the peak energy saved is:

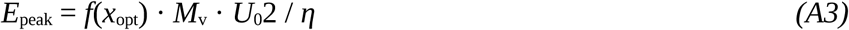

where *f*(*x*_opt_) = *g* · *x*_opt_ − ½(1 − 1/(1 + *x*_opt_)2) evaluates the dimensionless net mechanical energy balance at the optimal glide parameter *x*_opt_ = *k U*_0_ *T*_opt_ = (√*g* − 1) · *Φ*(*U*_0_) (equation 10). The first term *g* · *x*_opt_ = *P*_ss_ · *T*_opt_ · *η* / (*M*_v_ · *U*_0_2) represents the mechanical energy saved by not fluking over *T*_opt_. The second term ½(1 − 1/(1 + *x*_opt_)2) = Δ*KE* · *η* / (*M*_v_ · *U*_0_2) represents the mechanical work required to re-accelerate from *v*(*T*_opt_) back to *U*_0_. This is equation (6) of the main text. Substituting the expressions for *P*_ss_ (equation 5) and *T*_opt_ (equation 3), *E*_peak_ may equivalently be written as *P*_ss_ · *T*_opt_ · *η*_eff_, where the species-invariant efficiency factor *η*_eff_ = *f*(*x*_opt_) / (*g* · *x*_opt_) takes values 0.83, 0.79, and 0.68 at 1.5, 3.0, and 6.0 m s−1 respectively.

### Structure of *Φ*(*U*_0_)

The functional form of *Φ*(*U*_0_) follows from analytical optimization of the quadratic-drag burst-glide cycle under quasi-steady assumptions (Weihs, 1974). The specific linear approximation and its coefficients (*a* = 0.58, *b* = 0.046 s m−1) are derived and calibrated in this paper. The optimal glide duration *T*_opt_ = (1/*α*) · (√*g* − 1) · *Φ*(*U*_0_) is derived from the condition that *S*(*T*) is maximized under the Weihs (1974) burst-glide framework, from which the factor (√*g* − 1) emerges analytically as the primary driver of the optimum. The velocity-dependent correction *Φ*(*U*_0_) accounts for higher-order terms arising from the cubic scaling of active swimming power and the finite re-acceleration cost. A first-order Taylor expansion over the biologically relevant speed range 0.5 to 6.0 m s−1 yields *Φ*(*U*_0_) ≈ 0.58 + 0.046 *U*_0_, reducing *T*_opt_ to 60% of the zero-order Weihs prediction at 0.5 m s−1 and to 86% at 6.0 m s−1. The coefficients *a* = 0.58 and *b* = 0.046 s m−1 were calibrated to reproduce the optimal glide durations for *Tursiops truncatus* reported by Fish (1993). No additional species-specific fitting was performed.

### Structure of *Ψ*(*U*_0_)

The function *Ψ*(*U*_0_) and the associated maximum glide duration *T*_zero_ are introduced in this paper. The maximum glide duration *T*_zero_ is defined by *S*(*T*_zero_) = 0 for *T*_zero_ > 0. Expressing *T*_zero_ as a multiple of *T*_opt_ yields *Ψ*(*U*_0_) = *T*_zero_/*T*_opt_, approximated over the same speed range by *Ψ*(*U*_0_) ≈ 2.18 + 0.128 *U*_0_. The linear approximation is fitted to values of Ψ computed numerically from the profitability function (equation A1) at the three benchmark speeds [1.5, 3.0, 6.0] m s⁻¹. Fit residuals are below 0.01 across the full speed range. The monotonically increasing ratio reflects the increasing re-acceleration penalty at higher speeds, consistent with the cubic scaling of active swimming power on speed. Both *Φ* and *Ψ* are species-independent, as they depend only on the shared *g*(*U*) and the quasi-steady drag formulation. Sensitivity analysis confirms that moderate perturbations to the coefficients alter absolute *T*_opt_ and *T*_zero_ values but do not affect the algebraic cancellation in the speed retention expression, which holds unconditionally for any combination of these parameters.

## Acknowledgements

…

## Supplementary

**Figure S1.**
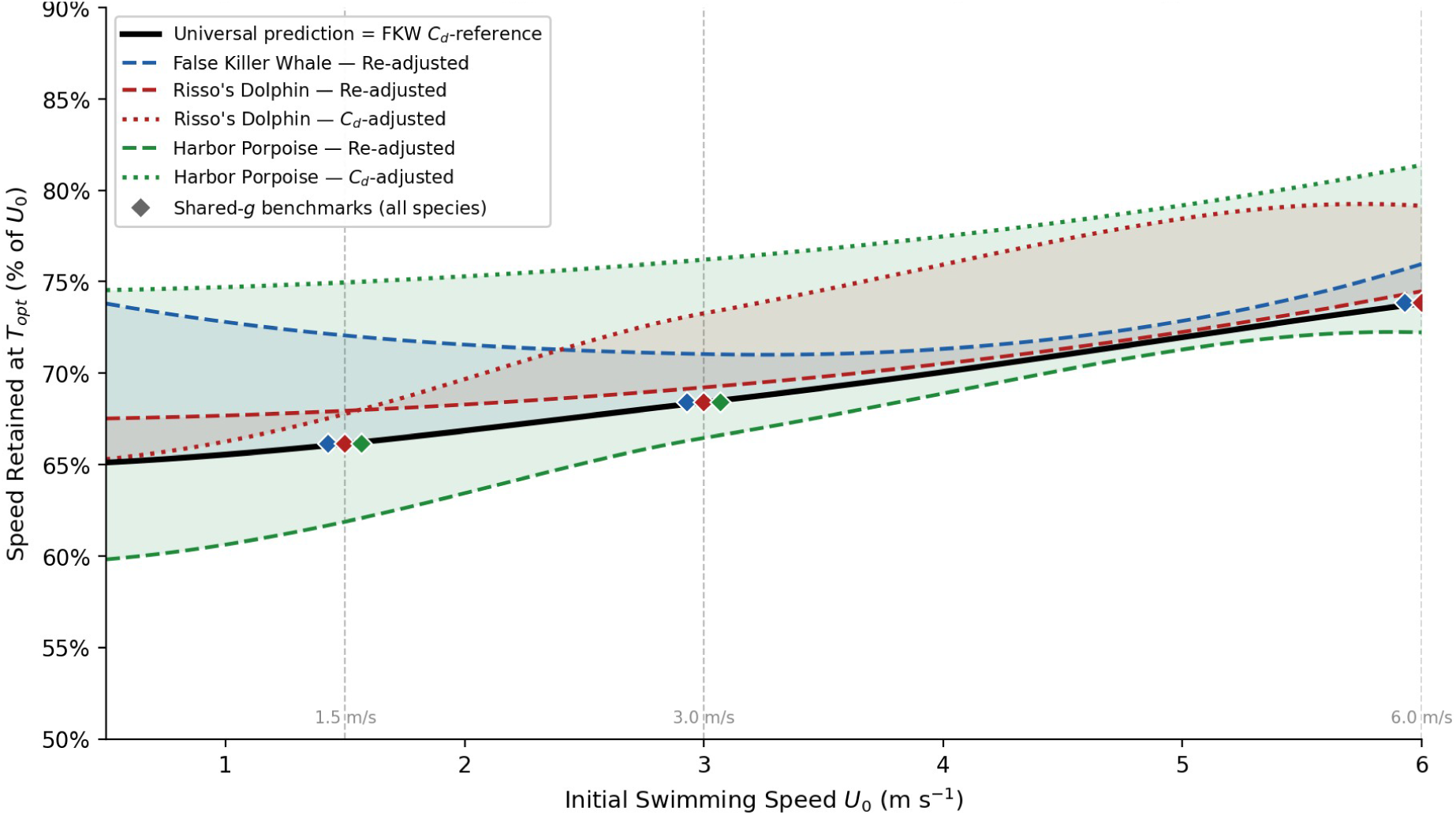
Speed retained at the optimal glide duration *T*_opt_ vs initial swimming speed *U*₀. The solid black line is the body-form-invariant model prediction under shared *g*(*U*) (equation 10). Diamond markers show benchmark values at 1.5, 3.0, and 6.0 m s⁻¹. All three species fall on the same curve, illustrating the body-form invariance of speed retention. For each species, dashed lines show the Re-adjusted scenario and dotted lines show the *C*_d_-correction scenario, two independent estimates of how the species’ drag ratio departs from the shared value. Shaded regions indicate the range spanned by the two scenarios, which narrows where the estimates coincide.

## References

1. Arranz, P., Benoit-Bird, K. J., Friedlaender, A. S., Hazen, E. L., and Goldbogen, J. A. (2019). Kinematics of Risso’s dolphins foraging dives in the Southern California Bight. Journal of Experimental Biology 222, jeb194225.

2. Bale, R., Hao, M., Bhalla, A. P. S., and Patankar, N. A. (2014). Energy efficiency and allometry of movement of swimming and flying animals. Proceedings of the National Academy of Sciences 111, 7517–7521.

3. Borazjani, I., and Sotiropoulos, F. (2008). Numerical investigation of the hydrodynamics of carangiform swimming in the transitional and inertial flow regimes. Journal of Experimental Biology 211, 1541–1558.

4. Brennen, C. E. (1982). A review of added mass and fluid inertial forces. Report CR 82.010, Naval Civil Engineering Laboratory, Port Hueneme, California.

5. Brès, G. A., Bose, S. T., Emory, M., Ham, F. E., Schmidt, O. T., Rigas, G., and Colonius, T. (2018). Large-eddy simulations of co-annular turbulent jet using a Voronoi-based mesh generation framework. In Proceedings of the AIAA/CEAS Aeroacoustics Conference, Atlanta, GA, USA, p. 3302.

6. Buckingham, E. (1914). On Physically Similar Systems; Illustrations of the Use of Dimensional Equations. Physical Review 4, 345–376.

7. Fish, F. E. (1993). Power output and propulsive efficiency of swimming bottlenose dolphins (Tursiops truncatus). Journal of Experimental Biology 185, 179–193.

8. Fish, F. E. (1998). Comparative kinematics and hydrodynamics of odontocete cetaceans. Journal of Experimental Biology 201, 2867–2877.

9. Fish, F. E., Legac, P., Williams, T. M., and Wei, T. (2014). Measurement of hydrodynamic force generation by swimming dolphins using bubble DPIV. Journal of Experimental Biology 217, 252–260.

10. Fish, F. E. and Lauder, G. V. (2006). Passive and active flow control by swimming fishes and mammals. Annual Review of Fluid Mechanics 38, 193–224.

11. Floryan, D., Van Buren, T. and Smits, A. J. (2017). Forces and energetics of intermittent swimming. Acta Mechanica Sinica 33, 725–732.

12. Gazzola, M., Argentina, M., and Mahadevan, L. (2014). Scaling macroscopic aquatic locomotion. Nature Physics 10, 758–761.

13. Gleiss, A. C., Wilson, R. P., and Shepard, E. L. C. (2011). Making overall dynamic body acceleration work: on the theory of acceleration as a proxy for energy expenditure. Methods in Ecology and Evolution 2, 23–33.

14. Glarou, M., Christiansen, F., Iwata, T., Basran, C. J., Ruppert, S. N. S., Sotiropoulou, D., Iversen, M. R., Akamatsu, T., Schnitzler, J. G., Siebert, U., and Rasmussen, M. H. (2025). Respiration rates and inferred mass-specific field metabolic rates decline with body size among five sympatric cetaceans. Journal of Comparative Physiology B 195, 659–675.

15. Gough, W. T., Segre, P. S., Bierlich, K. C., Cade, D. E., Potvin, J., Fish, F. E., et al. (2019). Scaling of oscillatory kinematics and Froude efficiency in baleen whales. Journal of Experimental Biology 222, jeb196287.

16. Kawai, S., and Larsson, J. (2012). Wall-modeling in large eddy simulation: Length scales, grid resolution, and accuracy. Physics of Fluids 24, 015105.

17. Li, G., Kolomenskiy, D., Liu, H., Godoy-Diana, R. and Thiria, B. (2023). Intermittent versus continuous swimming: An optimization tale. Physical Review Fluids 8, 013101.

18. Lamb, H. (1932). Hydrodynamics, 6th edn. Cambridge University Press, Cambridge.

19. Lighthill, M. J. (1971). Large-amplitude elongated-body theory of fish locomotion. Proceedings of the Royal Society of London B 179, 125–138.

20. Liu, A., Xing, C., Cao, W., Liu, B., Wang, S., Jin, J., Zhang, W., Li, K., Lu, Y., Hao, Y. and Cao, Y. (2025a). Bio-inspired intermittent locomotion: A novel energy-saving strategy for manta ray-inspired underwater vehicles. Ocean Engineering 341, 122425.

21. Liu, H., Priya, S. and James, R. D. (2025b). Comprehensive scaling laws across animals, microorganisms and plants. Proceedings of the Royal Society A 481, 20250515.

22. Miller, P. J. O., Johnson, M. P., Tyack, P. L., and Terray, E. A. (2004). Swimming gaits, passive drag and buoyancy of diving sperm whales. Journal of Experimental Biology 207, 1953–1967.

23. Noren, S. R., Williams, T. M., Ramirez, K., Boehm, J., Glenn, M., and Cornell, L. (2006). The physiology of diving in three odontocete cetacean species. Journal of Comparative Physiology B 176, 787–797.

24. Paoletti, P. and Mahadevan, L. (2014). Intermittent locomotion as an optimal control strategy. Proceedings of the Royal Society A 470, 20130535.

25. Pavlov, V., Vincent, C., Mikkelsen, B., Lebeau, J., Ridoux, V., and Siebert, U. (2021). Form, function, and divergence of a generic fin shape in small cetaceans. PLOS ONE 16, e0255464.

26. Rayner, J. M. (1985). Bounding and undulating flight in birds. Journal of Theoretical Biology 117, 47–77.

27. Rohr, J. J. and Fish, F. E. (2004). Strouhal numbers and optimization of swimming by odontocete cetaceans. Journal of Experimental Biology 207, 1633–1642.

28. Saadat, M., Fish, F. E., Domel, A. G., Di Santo, V., Lauder, G. V., and Haj-Hariri, H. (2017). On the rules for aquatic locomotion. Physical Review Fluids 2, 083102.

29. Sato, K., Watanuki, Y., Takahashi, A., Miller, P. J. O., Tanaka, H., Kawabe, R., et al. (2007). Stroke frequency, but not swimming speed, is related to body size in free-ranging seabirds, pinnipeds and cetaceans. Proceedings of the Royal Society B 274, 471–477.

30. Skandalis, D. A., Segre, P. S., Bahlman, J. W., Groom, D. J. E., Welch, K. C., Witt, C. C., McGuire, J. A., Dudley, R., Lentink, D., and Altshuler, D. L. (2017). The biomechanical origin of extreme wing allometry in hummingbirds. Nature Communications 8, 1047.

31. Schlichting, H. (1979). Boundary Layer Theory, 7th edn. McGraw-Hill, New York.

32. Schmidt-Nielsen, K. (1972). Locomotion: Energy cost of swimming, flying, and running. Science 177, 222–228.

33. Skrovan, R. C., Williams, T. M., Berry, P. S., Moore, P. W., and Davis, R. W. (1999). The diving physiology of bottlenose dolphins (Tursiops truncatus). II. Biomechanics and changes in buoyancy at depth. Journal of Experimental Biology 202, 2749–2761.

34. Watanabe, Y. Y., Sato, K., Watanuki, Y., Takahashi, A., Mitani, Y., Amano, M., et al. (2011). Scaling of swim speed in breath-hold divers. Journal of Animal Ecology 80, 57–68.

35. Webb, P. W. (1975). Hydrodynamics and energetics of fish propulsion. Bulletin of the Fisheries Research Board of Canada 190, 1–158.

36. Weihs, D. (1974). Energetic advantages of burst swimming of fish. Journal of Theoretical Biology 48, 215–229.

37. Williams, T. M. (1999). The evolution of cost efficient swimming in marine mammals. Philosophical Transactions of the Royal Society B 354, 193–201.

38. Williams, T. M., et al. (2000). Sink or swim: strategies for cost-efficient diving by marine mammals. Science 288, 133–136.

39. Williams, T. M., Fuiman, L. A., Kendall, T., Berry, P., Richter, B., Noren, S. R., et al. (2017). Exercise at depth alters bradycardia and incidence of cardiac anomalies in deep-diving marine mammals. Nature Communications 8, 14055.

40. Williams, T. M. (2022). Racing time: physiological rates and metabolic scaling in marine mammals. Integrative and Comparative Biology 62, 1439–1447.

41. Wisniewska, D. M., Johnson, M., Teilmann, J., Siebert, U., Galatius, A., Dietz, R., and Madsen, P. T. (2018). High rates of vessel noise disrupt foraging in wild harbour porpoises (Phocoena phocoena). Proceedings of the Royal Society B 285, 20172314.

42. Vreman, A. W. (2004). An eddy-viscosity subgrid-scale model for turbulent shear flow: algebraic theory and applications. Physics of Fluids 16, 3670–3681.

43. Zhu, Y., Kang, L., Ma, J., Tian, F. and Fan, D. (2025). Intermittent swimmers optimise energy expenditure with flick-to-flick motor control. Journal of Fluid Mechanics 1006, paper 45.

